# Upstream ORFs regulate translation of the Arabidopsis ZIF2 transporter and affect ER stress tolerance

**DOI:** 10.64898/2026.02.12.705568

**Authors:** Esther Novo-Uzal, Dóra Szakonyi, Paula Duque

## Abstract

Translational regulation mediated by upstream open reading frames (uORFs) is an important mechanism controlling gene expression in eukaryotes, yet the functional characterization of uORFs in plant transporter genes remains limited. ZINC-INDUCED FACILITATOR 2 (ZIF2) is an *Arabidopsis thaliana* tonoplast-localized zinc transporter whose expression is regulated at the translational level via alternative splicing of its 5’ untranslated region (UTR), but the role of additional *cis*-regulatory elements within this region has remained unexplored. Here, we characterize the functional significance of three uORFs present in the *ZIF2* 5’UTR and elucidate their contribution to translational control and stress responses. We demonstrate that two of the three uORFs (uORF2 and uORF3) act as negative regulators of *ZIF2* translation. uORF2 is constitutively translated and plays the primary role in uORF-mediated translational inhibition, whereas uORF3 functions as a fail-safe mechanism that represses translation only when uORF2 is inactive. We further show that uORF2-mediated repression depends on the amino acid sequence of the encoded peptide and identify a short motif critical for its function. Importantly, uORF-mediated regulation of *ZIF2* is physiologically relevant *in planta*: plants carrying mutations that abolish uORF2 translation accumulate higher levels of ZIF2 protein and display increased tolerance to endoplasmic reticulum stress. Together, our findings reveal a uORF-based fail-safe regulatory mechanism that modulates *ZIF2* expression, contributes to plant stress adaptation, and is consistent with translational inhibition mediated by peptide-dependent ribosome stalling.

## INTRODUCTION

The 5’ untranslated region (UTR) of the mRNA contains different regulatory elements, such as protein binding sites and upstream Open Reading Frames (uORFs), that are key to translational regulation. Translation initiation represents not only the first step of protein synthesis but also the central hub of translational regulation, as it determines the overall rate of protein production. uORFs are protein-coding regions in mRNAs that lie upstream of the main ORF (mORF), *i*.*e*. in the 5’ UTR of the mRNA. uORFs are known to be able to affect translation of the mORF, usually reducing the levels of protein production due to a decrease in translation efficiency (Calvo et al 2009, Zhang et al 2020).

Around 20-50% of eukaryotic genes contain uORFs, and in *Arabidopsis thaliana* these genes have been estimated to be 35%, with about half harboring multiple uORFs. Other plant species have similar fractions of transcripts with uORFs (von Arnim et al 2014). The distribution of uORFs is strongly associated with gene function. mRNAs that are highly expressed, including many housekeeping genes, typically possess short 5’ UTRs lacking uORFs. In contrast, transcripts expressed at lower levels, such as those encoding transcription factors and kinases, often contain longer 5’ UTRs enriched in uORFs (Kim et al 2007). These findings indicate that the presence or absence of uORFs is likely adaptive. Nevertheless, only a limited number of uORFs have been directly investigated for their functional relevance at the whole-plant level (Ribone et al 2017, Aibara et al 2018, Reis et al 2020). Therefore, uORF control of plant developmental mechanisms is poorly established, even though over 40% of mRNAs for protein kinases and transcription factors harbor uORFs (Kim et al 2007).

ZIF2 (Zinc-Induced Facilitator 2) is an *A. thaliana* membrane transporter belonging to the Major Facilitator Superfamily (Niño-González et al 2019) that localizes at the tonoplast of root cortical cells and mediates vacuolar compartmentalization of zinc ions. This prevents zinc from reaching the aerial part of the plant where it would cause detrimental effects, thereby conferring tolerance to heavy metal (Remy et al 2014). This MFS gene shows post-transcriptional regulation, and it undergoes alternative splicing, generating two variants that differ only in the 5’UTR region due to an intron retention event (Remy et al 2014).

Information regarding the function of uORFs in translation of transporter encoding genes is scarce. However, it has been reported that the *A. thaliana* boron (B) transporters NIP5;1 and BOR1 are regulated by a uORF (Tanaka et al 2016, Aibara et al 2018). Recently, the phosphate transporter PHO1 mRNA was reported to harbor a uORF able to repress translation and modulate phosphate levels in the shoot (Reis et al 2020). NIP5;1 encodes a boric acid channel required for normal growth under low B conditions. However, high levels of B promote the stalling of the ribosome at the uORF thus suppressing translation and mRNA degradation as well (Tanaka et al 2016, 2024). Stalling of the ribosome is one of the most common mechanisms by which uORFs prevent translation of the mORF. The peptide encoded by the uORF can stall the ribosome as a nascent peptide while located in the ribosome exit tunnel, thus blocking the progression of upstream ribosomes or suppressing reinitiation (Hanfrey et al 2002, Rahmani et al 2009). In other cases, the uORF peptide may exert its function after it has been released from the ribosome, thus acting in *trans*. Only two such cases have been described in plants (Chang et al 2000, Combier et al 2008).

The function of uORFs can depend on their specific sequences. However, the majority of uORFs shows sequence-independent functionality, which in turn relies on the context surrounding the uORF start codon or on the location of the uORF. On the other hand, sequence-dependent uORFs produce nascent peptides that may stall the ribosomes. These usually encode amino acid sequences with a high degree of conservation and are named as Conserved Peptide uORFs (CPuORFs) (van der Horst et al 2019, Zhang et al 2020).

Stress triggers a global reduction in protein synthesis, which allows the conservation of resources and enables extensive reprogramming of gene expression. Concurrently, a subset of mRNAs undergoes translation through uORF-dependent regulatory mechanisms, contributing to adaptive stress responses (Young & Wek 2016). The similar prevalence of uORFs in both translationally repressed and activated mRNAs during stress underscores that uORF-specific characteristics, rather than abundance, dictate their regulatory potential (Young & Wek 2016). Proteins produced from these selectively translated mRNAs fulfill diverse roles in stress alleviation.

uORFs have been recently used to engineer plants by classical transgenic technologies and cutting-edge genome editing in order to obtain crops resistant to biotic or abiotic stress without fitness costs (Xu et al 2017, Zhang et al 2018). This is particularly important because, usually, enhanced resistance obtained through ectopic expression of plant defense genes is associated with substantial penalties to fitness (Gurr et al 2005), making the resulting products undesirable for agricultural applications. Remarkably, uORF-engineered rice plants resist a spectrum of pathogens without compromising plant fitness in laboratory conditions or even in the field (Xu et al 2017), and lettuce plants genome edited in uORFs are able to resist oxidative stress better and accumulate more vitamin C (Zhang et al 2018). This strategy of uORF genome editing has been also expanded to enhance crop traits, such as rice grain shape (Yang et al 2025) or strawberry sugar content (Xing et al 2020). Therefore, uORF edition hence holds much promise to improved nutrition and better agricultural practices.

Here, we reveal the functional significance of the three uORFs present in the *A. thaliana ZIF2* gene in translational regulation and plant stress responses. Specifically, we elucidate the molecular mechanism governing *ZIF2* uORF function, using *in vitro* and *in vivo* approaches to show how these elements inhibit mRNA translation. We further establish the physiological relevance of uORF-mediated regulation of *ZIF2* translation at the whole-plant level in response to stress.

## MATERIALS AND METHODS

### Plant materials

The *A. thaliana* Columbia ecotype (Col-0) was used as the wild type in all experiments. The *zif2* knockout homozygous line (SALK_037804) was originally isolated in the Duque lab (Remy et al, 2014). The *upf1-5* and *upf3-1* mutants were kindly provided by M. Kalyna (University of Natural Resources and Life Sciences, Vienna, Austria). Plants were grown in growth chambers under long-day photoperiod: 16-h light/8-h dark at 21–24°C (100–120 µmol m^-2^ s^-1^ light intensity)/18°C (dark) and 60–65% relative humidity (RH). For phenotypical assays, unless otherwise specified, seeds were surface-sterilized and plated on MS medium (MS basal salt mix, 0.01% [w/v] myo-inositol, 0.05% [w/v] MES, pH 5.7, 0.8% [w/v] agar), and stratified for 3 days at 4°C in the dark before the transfer of the plates to a growth cabinet under long-day conditions (22°C light 30% RH/18°C dark 70% RH, 100 µmol m^-2^ s^-1^ light intensity). For DTT treatment, seeds were germinated in MS plates containing 2.5 mM DTT and grown under long-day conditions for 14 days.

### Preparation of DNA constructs for protoplast transfection

Constructs were made in a pUC18-derived vector expressing firefly luciferase (F-LUC) under the 35S promoter and NOS terminator (Martinho et al 2015). Constructs previously generated (Remy et al 2014) were modified in the 5’UTR by site-directed mutagenesis (Nyztech) in order to change uORFs context and abolish start and/or stop codons. For trans effect experiments, the vector pGL4.70, kindly given by K. Berendzen (University of Tuebingen, Germany), harboring firefly (*Photinus pyralis*) and sea pansy (*Renilla reniformis*) luciferases under the control of 35S promoter was used. Constructs derived from each species harboring the *ZIF2* 5’UTR with only one uORF active (achieved by disrupting the other uORF’s start codon) were compared with the vector harboring all uORFs mutated.

### Isolation and transfection of Arabidopsis Protoplasts

Plasmid DNA for protoplast transfection was purified from CsCl gradients and stained with a RedSafe Nucleic Acid Staining Solution (ChemBio; 1:500). Protoplasts were isolated from mature fully expanded leaves from 5-week-old plants as described (Confraria & Baena-Gonzalez 2016). For qRT-PCR experiments and LUC activity assays 2×10^5^ or 2×10^4^ protoplasts were transfected, respectively, using a ratio of 1 µg plasmid DNA per 1×10^3^ transfected protoplasts. In the LUC activity assays, 9 µg of construct were used, in combination with 1 µg of 35S:GUS (Martinho et al 2015) as transfection control. In the assays for the relative quantification of *LUC* transcript levels, the same components were scaled up 10-fold, to maintain the ratio of plasmid DNA per transfected protoplasts. mER7 plasmid was used as control DNA (Kovtun et al 1998). After transfection, protoplasts were incubated overnight under light (15 µE; 25°C). On the following day, protoplasts were harvested by centrifugation at 100 *g* for 3 min, flash-frozen in dry ice, and used for RNA extraction or for luciferase and β-glucuronidase analyses (Yoo et al 2007). To compare LUC activity among different samples, activity values were normalized to β-glucuronidase activity derived from the co-transfected 35S:GUS reporter.

### Generation of transgenic Arabidopsis lines

Constructs harboring the *ZIF2* native 5’UTR or the 5’UTR carrying mutations in uORF2 and uORF3, either individually or in combination, were generated. We used the previously described 5’UTR:ZIF2-YPF construct driven by the strong constitutive 35S promoter as a backbone to introduce the corresponding mutations in the 5’UTR by site-directed mutagenesis (Remy et al 2014). The different constructs were then used to transform the *Arabidopsis zif2* mutant by the floral dip method (Clough & Bent 1998) using *Agrobacterium tumefaciens* strain GV3101.

### RNA extraction and qRT-PCR analyses

Total RNA was extracted from *Arabidopsis* seedlings using the innuPREP Plant RNA kit (Analytik Jena BioSolutions). All RNA samples were treated with DNase I (Promega), and cDNA was synthesized using SuperScript III reverse transcriptase (Invitrogen) and oligo(dT) 18 primers. qRT-PCR was performed using an ABI QuantStudio 384-well instrument (Applied Biosystems) and Luminaris Color HiGreen qPCR master mix High ROX (Thermo Scientific) with 2.5 µl of cDNA (diluted 1:10 for plant RNA or 1:5 for protoplasts RNA) per 10-µl reaction volume containing 300 nM of each specific primer (Supplemental Table 1). Relative quantification was assessed using the 2ΔΔCt method (Livak & Schmittgen 2001) and GUS as a reference gene for protoplasts experiments and UBQ or SAND for plant experiments. Statistical differences between the average relative expression values of individual samples were inferred using Student’s t-test. Statistical analyses were conducted using GraphPad Prism (v. 9.1.1, GraphPad Software, Boston, MA, USA).

### Protein extraction and western blotting

Roots from 10-day-old *Arabidopsis* seedlings were ground with a mortar and pestle under liquid nitrogen. Total protein was extracted (3:1, v:w) in an extraction buffer containing 50 mM Tris-HCl pH 8.0, 150 mM NaCl, 1 mM EDTA, 0.5% Triton X-100 (Sigma-Aldrich), 0.5% sodium deoxycholate, 0.1% SDS and Complete Protease Inhibitor Cocktail (Roche). The extract was centrifuged at 12 000 g for 30 min at 4º C and the supernatant was mixed with Laemmli buffer. Proteins were resolved on 10% SDS–polyacrylamide gels and then transferred to PVDF membranes (Immobilon-P, Millipore) and blocked with 5% non-fat dry milk for 2 h. The membranes were probed overnight at 4 C with anti-GFP primary antibodies (Roche, 11814460001; 1:1000) and then with anti-mouse peroxidase-conjugated secondary antibodies (Jackson ImmunoResearch, 115-035-146; 1:20000) for 2 h at room temperature. All antibodies were diluted in TBS buffer (25 mM Tris-HCl [pH 7.4] and 137 mM NaCl) supplemented with 1% non-fat dry milk. After incubation with the antibodies, the membranes were washed with TBS containing 0.05% Tween 20 (Sigma-Aldrich) for 40 min. The peroxidase activity associated with the membrane was visualized by enhanced chemiluminescence. The intensity of the protein bands was quantified using ImageJ software, normalizing protein levels to that of the Rubisco large subunit visualized in membranes stained with 0.1% Ponceau S (Sigma-Aldrich) in 5% acetic acid. Statistical differences between the average protein levels of each sample were inferred using Student’s t-test.

In the case of *Arabidopsis* mesophyll protoplasts, protein was extracted directly with Laemmli buffer by incubating at 95º for 10 min and resolved in 7.5% polyacrylamide gels. For western blot, we followed the protocol described above, using C-LUC primary antibody (Santa Cruz Biotechnology, sc-74548, 1:100).

## RESULTS

### *ZIF2* 5’UTR contains uORFs that inhibit translation

In a previous work, we demonstrated that the ZIF2 5’UTR was crucial for post-transcriptional regulation of this gene (Remy et al 2014). To gain further insight into translational regulation of the *ZIF2* gene, we investigated the putative role of *cis* elements in its 5’UTR, finding that *ZIF2* mRNA contains three upstream Open Reading Frames (uORF) in the 5’UTR (Fig. 1). uORF1 is the longest, spanning 39 amino acids; uORF2 is 14 amino acids long and uORF3, the closest to the mORF, is the shortest with just 9 amino acids. uORF1 overlaps uORF2 and uORF3 sequences, while both uORF2 and uORF3 are in frame (Fig. 1).

**Figure 1.**
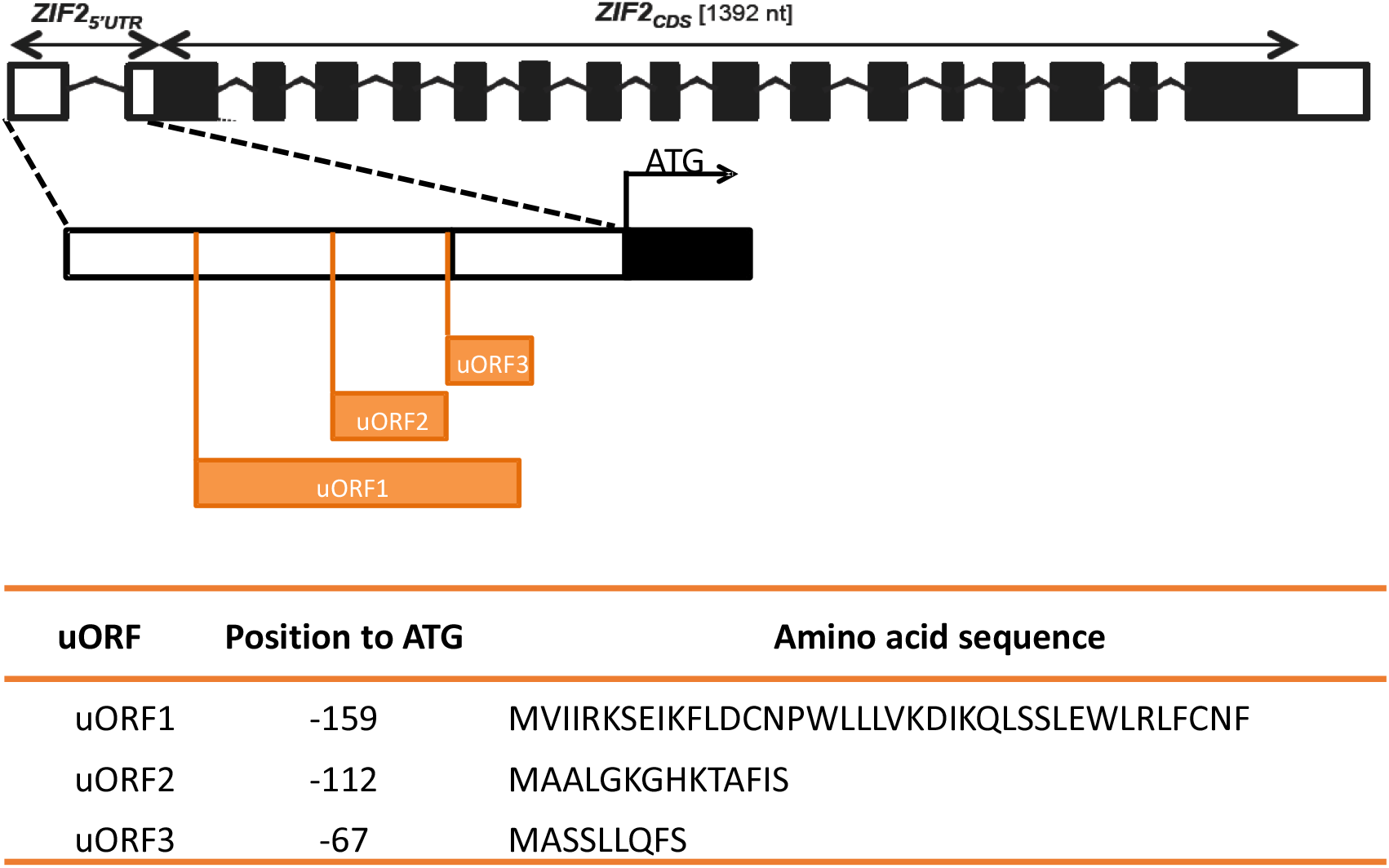
*ZIF2* gene contains three uORFs in its 5’UTR. Schematic diagram of the *ZIF2* 5’UTR showing the location of the different uORFs in *ZIF2* mRNA, and peptides encoded by the uORFs.

Our first objective was to assess whether the uORFs in *ZIF2* 5’UTR affect translation of the main ORF. To this end, we used site-directed mutagenesis to disrupt the start codon AUG (mutated to UUG) of the *ZIF2* uORFs and fused the 5’UTR to the luciferase (LUC) reporter gene for transient expression assays *Arabidopsis* mesophyll protoplasts (Fig. 2A). Simultaneous disruption of all uORFs caused a marked increase in LUC activity, indicating that at least one uORF is repressing translation of the main ORF (mORF) (Fig. 2C). To evaluate to which extent each individual uORF contributes to the translational inhibition, we disrupted the AUG start codons individually. When we mutated the AUG of uORF1, no change in LUC activity was detected, compared to WT 5’UTR. When mutating the start codon of both uORF2 and uORF3, therefore uORF1 being the only one putatively expressed, an increase in LUC activity, compared to mutating all uORFS was detected. These two results suggest that uORF1 does not inhibit mORF translation. On the contrary, when we mutated uORF2 AUG, LUC activity increased, but leaving uORF2 as the only uORF in *ZIF2* 5’UTR did not change LUC activity, concluding that uORF2 represses mORF translation. In the case of uORF3, we obtained apparent contradictory results, given that LUC activity did not change when mutating uORF3 start codon or when uORF3 is the only uORF expressed in the *ZIF2* 5’UTR. However, when taking a closer look to the position of the two uORFs in the 5’UTR, uORF3 is just after the stop codon of uORF2, implying that the 80S ribosome cannot resume uORF3 translation following the dissociation of ribosome subunits after uORF2 stop codon. In light of these results, we conclude that uORF2 is the main player inhibiting translation, whereas uORF1 does not affect translation (Fig. 2C). Moreover, we hypothesize that uORF3 may act as a fail-safe mechanism when uORF2 is not functional.

**Figure 2.**
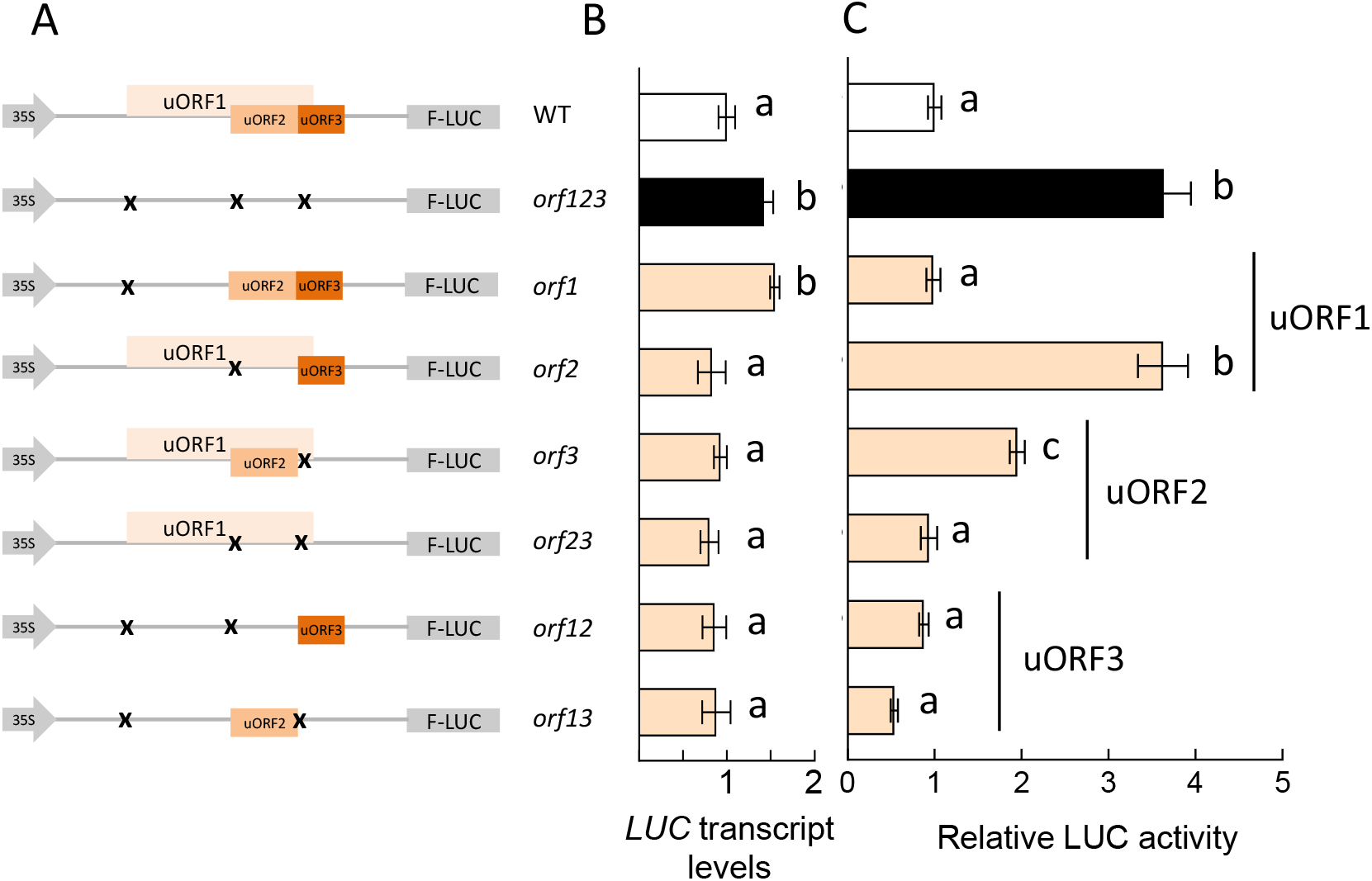
Effect of *ZIF2* uORFs in translation. **A.** Scheme of the constructs generated by site-directed mutagenesis. Orange boxes represent uORFs, while mutation of each start codon is indicated with an x. **B**. *LUC* transcript levels in different constructs carrying *ZIF2* uORF with mutated start codons. **C**. Effect on luciferase activity of *ZIF2* uORF start codons disruption. Average ± SE; *n* = 12. Different letters indicate statistically significant differences (ANOVA). The number after “*orf”* indicates the ORF(s) in which the AUG was mutated.

### uORF2 and uORF3 are translated

Our next goal was to investigate whether *ZIF2* uORFs are indeed translated and the corresponding peptides synthesized. *In silico* analysis of the start codon context indicated that translation of uORF2 and uORF3 is likely, and published data supported our hypothesis, at least for uORF2 (Hu et al 2016). To experimentally address this question, stop codons of *ZIF2* uORFs were independently mutated. As uORF1 is not in frame with the LUC start codon, we mutated the two stop codons in frame with the uORF1 AUG found upstream of the LUC start codon, using a construct in which both uORF2 and uORF3 start codons were disrupted (*orf23*). If uORF1 were translated, it would overlap the frame of LUC, therefore producing no protein and no LUC activity. On the contrary, results (Fig. 3A) showed that mutating uORF1 stop codons showed no change in LUC activity compared to its background (*orf23*), demonstrating that uORF1 is not translated and thereby explaining why uORF1 does not affect mORF translation (Fig. 2C).

**Figure 3.**
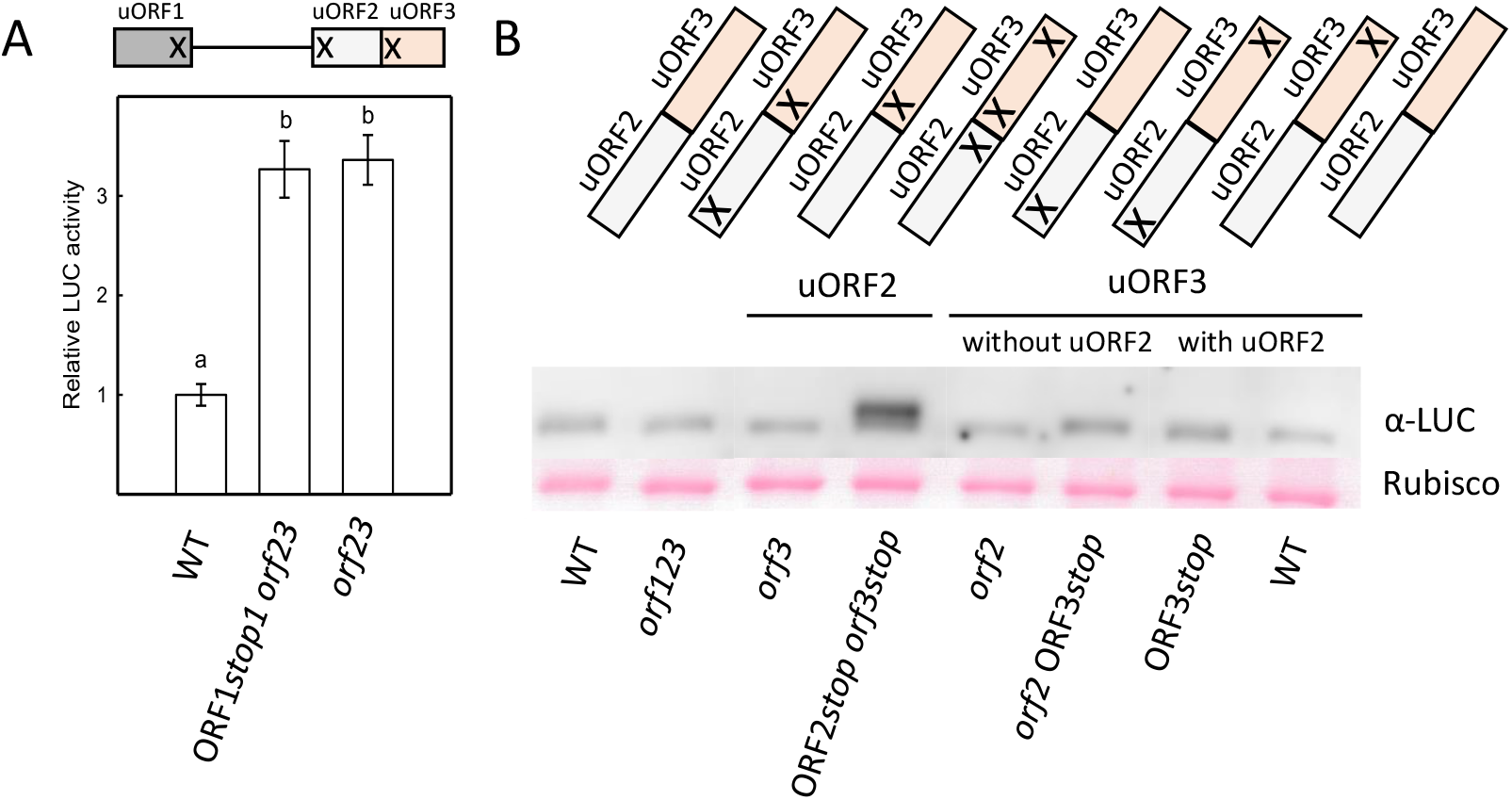
Translation of *ZIF2* uORFs. **A.** A construct in which stop codons of uORF1 were disrupted, as well as uORF2 and uORF3 start codons (*orf23*), was generated and used to transfect *Arabidopsis* protoplasts with the aim of assessing uORF1 translation. Different letters indicate statistically significant differences (ANOVA). Bars indicate average of 4 independent experiments ± SE. **B**. Western blot analysis using anti-luciferase (α-LUC) antibodies (upper panel) and Ponceau staining to detect RuBisCO as a loading control (lower panel). Protoplasts were transfected with different constructs, in which the stop codons of *ZIF2* uORF2 and uORF3 were mutated. Cells were then disrupted and subjected to denaturing separation in polyacrylamide gels (SDS-PAGE) followed by α-LUC blotting. A higher band indicates that the uORF is being translated because a fusion protein is produced (portion of 5’UTR starting at AUG of uORF + LUC).

On the other hand, uORF2 and uORF3 are in frame with LUC start codon. If those uORFs were translated, in absence of a stop codon, the ribosome would continue translating until the stop codon of LUC (the mORF, in this case), producing a larger protein than the original LUC, traceable with α-LUC antibodies by western blot analysis. The results obtained are depicted in Figure 3B and clearly show that both uORF2 and uORF3 are translated. Interestingly, uORF3 is only translated in the absence of uORF2, i.e., when uORF2 start codon is disrupted. When the uORF2 start codon is functional, no production of the uORF3-LUC protein is detected (Fig. 3B). These results are in agreement with the data obtained with the LUC activity experiments shown in Fig. 2C, thus unequivocally demonstrating that, at least in isolated *Arabidopsis* protoplasts, uORF3 acts as a fail-safe mechanism, being translated to inhibit translation when uORF2 happens not to be translated. This fine-tuning of translational regulation may have a crucial role in the function of ZIF2 and disclose important novel insight into the mechanisms governing plant responses.

### Molecular mechanisms underlying *ZIF2* uORFs function

Selection of the translation start site relies largely on the nucleotide context surrounding the start codon. The optimal sequence, named the Kozak consensus sequence, is GCC(A/G)CC<u>AUG</u>G, with the most important residues being the purines in the -3 and +4 positions and the underlined initiation codon (Kozak et al 1987). Poor uORF start codon context has also been associated with mRNAs that are preferentially translated, whereas mRNAs that are repressed under stress conditions typically contain an uORF with a strong Kozak consensus sequence (Baird et al 2014). These findings suggest that the start codon’s context plays a significant role in uORF-mediated translation regulation. To assess the importance of the AUG context for *ZIF2* uORF function, we used site-directed mutagenesis to alter the nucleotides in positions +4 and -3 (reported to determine the AUG context’s strength) to create weaker contexts in the start codons of uORF2 (changing UCC**AUG**G to UUC**AUG**U) and uORF3 (UGA**AUG**G to UGU**AUG**U). The generated constructs were transfected into isolated *Arabidopsis* protoplasts and LUC activity measured together with the non-mutated controls (Fig. S1). The results showed that LUC activity did not change when introducing mutations in the Kozak sequence, showing that the AUG context is not important for either uORF2-or uORF3-mediated inhibition of mORF translation (Fig S1).

In order to disclose the mode of action of *ZIF2* uORF2 and uORF3, we first investigated whether the *ZIF2* mRNA was targeted to Nonsense-Mediated Decay (NMD), given that the uORF stop codon can be recognized as premature and trigger NMD. To do so, *ZIF2* expression was detected by quantitative RT-PCR in *lba1* and *upf3-1* seedlings, two mutant lines for UPF1 and UPF3, two factors with a prominent role in NMD. The results showed that *ZIF2* expression levels were similar to *Actin*, used as a negative control, but clearly different to *SMG7*, an established NMD target (Lorenzo-Orts et al 2019), whose expression is therefore enhanced in the *lba1* and *upf3-1* mutants (Fig. 4A). Thus, the *ZIF2* mRNA does not appear to constitute an NMD target and this mRNA decay mechanism is likely not involved in the observed reduction of mORF protein expression (Fig. 2).

**Figure 4.**
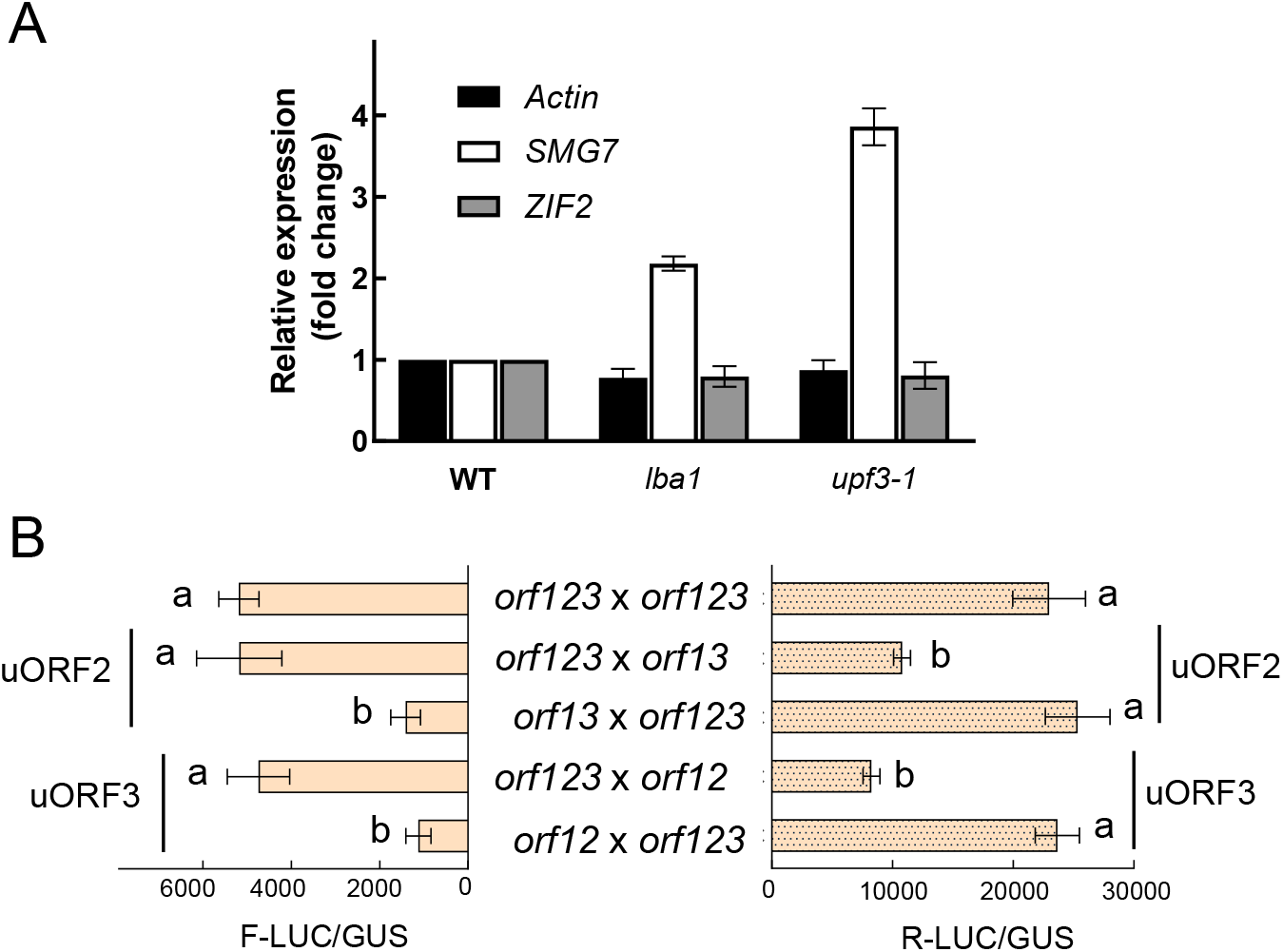
*ZIF2* uORFs mode of action. **A.** *ZIF2* is not a target of Nonsense-Mediated Decay (NMD). mRNA levels were quantified by RT-qPCR in WT (Col-0) and NMD mutants (*lba1* and *upf3-1*) 7 days after sowing, using *SAND* expression to normalize the results. *ZIF2* expression levels were compared to *Actin* (negative control) and *SMG7* (NMD target). **B**. uORF2 and uORF3 do not act in *trans*. Co-transfection of *Arabidopsis* protoplasts with LUC vectors of different origins, firefly (*Photinus pyralis*, F-LUC) and sea pansy (*Renilla reniformis*, R-LUC), expressing proteins using different LUC substrates, allowed distinguishing the origin of LUC activity. Constructs derived from each species harboring the *ZIF2* 5’UTR with only one uORF active were compared with the vector harboring all uORFs mutated. Bars indicate average of 3 independent experiments ± SD. Different letters indicate statistically significant differences (ANOVA).

Although infrequent, some uORF-encoded peptides can inhibit translation in a *trans* mode (Combier et al 2008). To test this hypothesis, we took advantage of two LUC vectors of different origins, one from firefly and the other from Renilla (Fig. 4B). Considering that the substrates for each LUC are different, we could determine from which vector LUC activity was arising after cotransfecting protoplasts with both vectors. Results show no decrease in LUC activity when the no uORFs vector was incubated with a vector expressing only either uORF2 or uORF3, meaning that uORF2 and uORF3 do not act in *trans*.

To address whether uORF2 and uORF3 functionality depends on their specific amino acid sequence, a frame shift in the nucleotide sequence of the two *ZIF2* uORFs that are translated (uORF2 and uORF3) was performed to change the amino acid sequence while disturbing minimally the nucleotide sequence. The frame shift was introduced after the uORF’s AUG to maintain the start codon, and the frame was readjusted just before the uORF’s stop codon, so the number of nucleotides and amino acids remained unchanged. In the case of uORF2, the introduction of one nucleotide after the start codon creates a new stop codon in frame. To avoid this, two nucleotides were introduced after the AUG. Protoplasts were then transfected with these constructs and LUC activity determined. Strikingly, the frame shift introduced in uORF2 increased LUC activity by four fold compared to its control (*orf3*, Fig. 5C), while no change in LUC activity was observed when the frame shift was introduced in uORF3. These results indicate that uORF2 represses translation of the mORF in a peptide sequence-dependent manner, while the amino acid sequence of the encoded peptide is not important for uORF3 function. To the best of our knowledge, this is the first time that peptide sequence dependence has been described for a non-conserved plant uORF.

**Figure 5.**
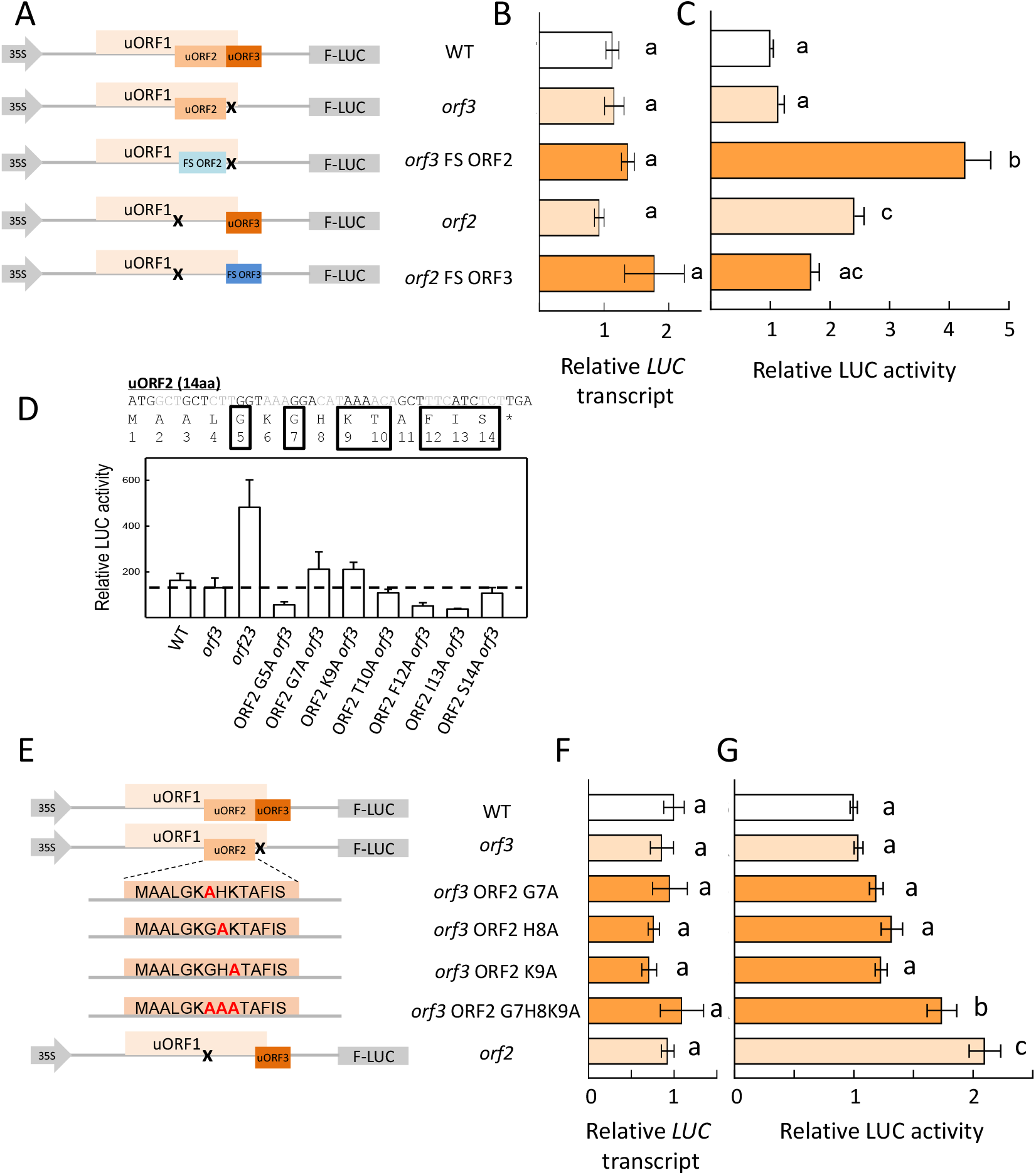
uORF2 inhibitory function depends on its encoded amino acid sequence. **A.** Scheme of the constructs generated by site-directed mutagenesis. Orange boxes represent uORFs, while blue boxes indicate the uORF carrying the frame shift (FS). Mutation of each start codon is indicated with an x. **B**. Relative *LUC* transcript levels of constructs represented in A after protoplast transfection. **C**. Effect of a FS in uORF2 and uORF3 sequence on LUC activity in arabidopsis protoplasts. FS experiments for uORF2 and uORF3 were performed in the absence of the uORF3 AUG (*orf3*) and the uORF2 AUG (*orf2*), respectively. **D**. Ala scanning effect on LUC activity in Arabidopsis protoplasts. **E**. Scheme of the constructs used for Ala scanning, where orange boxes represent uORFs. The mutated amino acids are colored in red. **F**. Relative LUC transcript leves of constructs in E, after protoplast transfection. **G**. Effect of G7H8K9 amino acid substitution to ala on LUC activity. Different letters indicate statistically significant differences (ANOVA). Bars indicate average of 7 independent experiments ± SE.

In order to identify the amino acids critical for uORF2-mediated translation inhibition, we performed an alanine scanning, by mutating several residues (marked with a black box) to Ala and see the mutation effect on translation. Results showed (Fig 5D) that only the changes in positions G7 and K9 slightly increased the LUC activity compared to its background (*orf3*). A mutant G7H8K9 showed a significant increase in LUC activity, pointing to a critical importance of these three residues. However, the LUC activity achieved in the mutant G7H8K9 is reduced compared to mutating uORF2 start codon alone (Fig. 5G), suggesting that another amino acids could minorly contribute to uORF2-mediated inhibition of translation.

These data, together with to the fact that *ZIF2* is not an NMD target, point to stalling of the ribosome as the mode of action for uORF2. In plants, stalling of the ribosome has been described as the most common mechanism by which uORFs inhibit the expression of the mORF (Tanaka et al 2016, Hayashi et al 2017, Ribone et al 2017). The ribosome stalling while translating the uORF prevents the ribosome from reaching the mORF and therefore inhibiting its translation.

### *ZIF2* uORFs inhibit translation *in planta*

We showed that *ZIF2* uORFs are able to repress translation in *Arabidopsis* mesophyll protoplasts *in vivo*, so we next aimed to test their role *in planta*. For this, we generated constructs with *ZIF2* fused to YFP with different versions of the 5’UTR, i.e., WT 5’UTR, and 5’UTR with mutations of start codon of uORF2, uORF3 or a combination of both uORFs. We generated different transgenic lines in the KO *zif2* background (Remy et al 2014), with different levels of transgene expression. Then, we identified the ZIF2-YFP protein with anti-GFP antibodies, and considering the different levels of *ZIF2* transcript, we calculated the efficiency of translation, dividing the quantified protein by the mRNA levels. Results show that the presence of uORF2 inhibits translation of *ZIF2* and this inhibition was greater when both uORF2 and uORF3 were combined (Fig. 6), mirroring the results obtained with protoplasts (Fig. 2C). This way, we were able to ascertain the functional relevance of *ZIF2* uORF-mediated translational regulation at the organism level.

**Figure 6.**
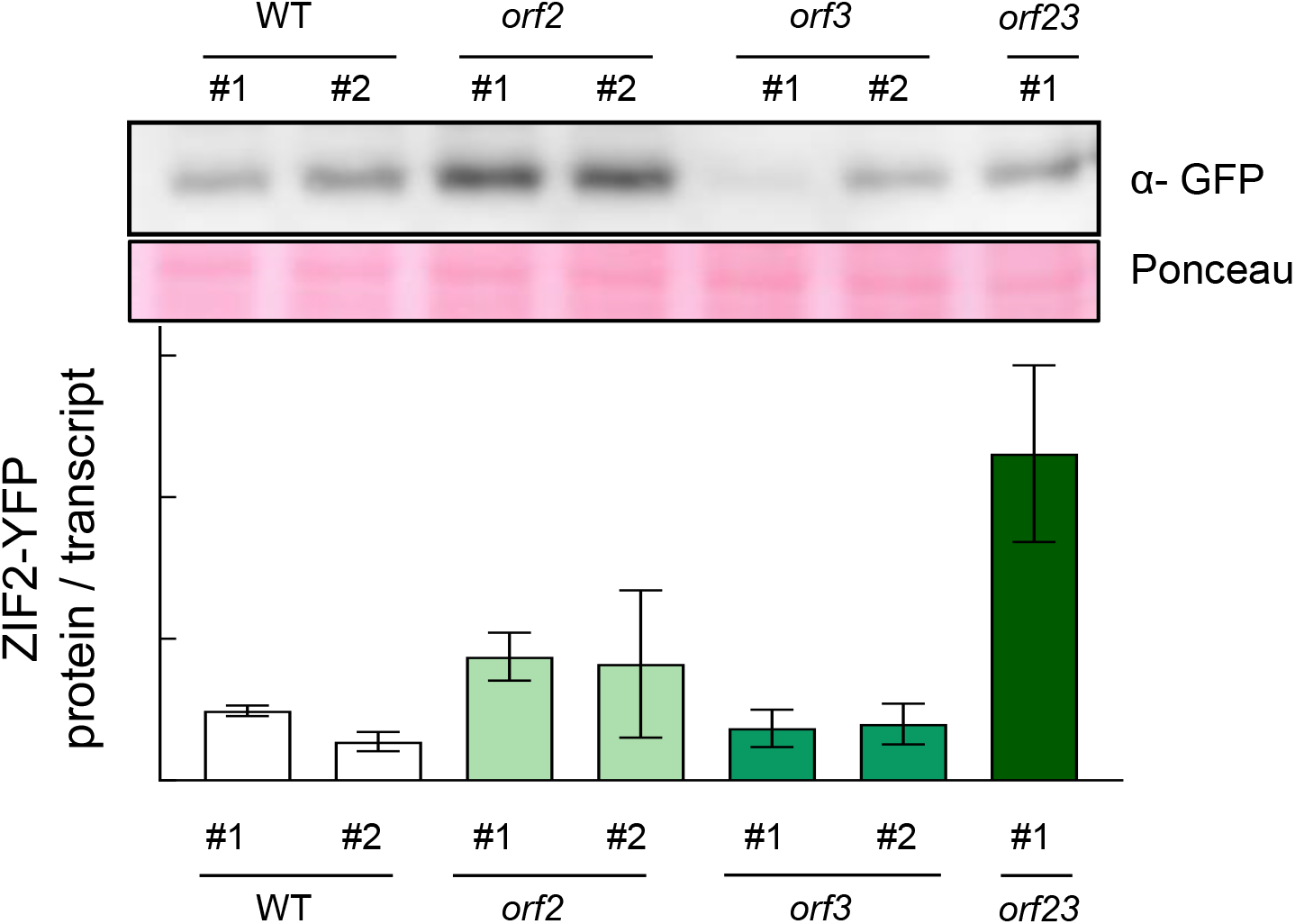
*ZIF2* uORFs inhibit translation in planta. *Arabidopsis ZIF2* knockout mutants were transformed with constructs harboring the native *ZIF2* 5’UTR or the 5’UTR with uORF2 and uORF3 mutated, individually or in combination. Seedlings were grown for 7 days, and after protein extraction, western blot analysis was performed using α-GFP antibodies and Ponceau staining to detect RuBisCO as a loading control. Protein bands were quantified with ImageJ and normalized against *ZIF2* transcript levels (evaluated by quantitative RT-PCR) to obtain the ZIF2-YFP protein/transcript ratio for each transgenic line. Bars indicate average of 3 independent experiments ± SD.

**Figure 7.**
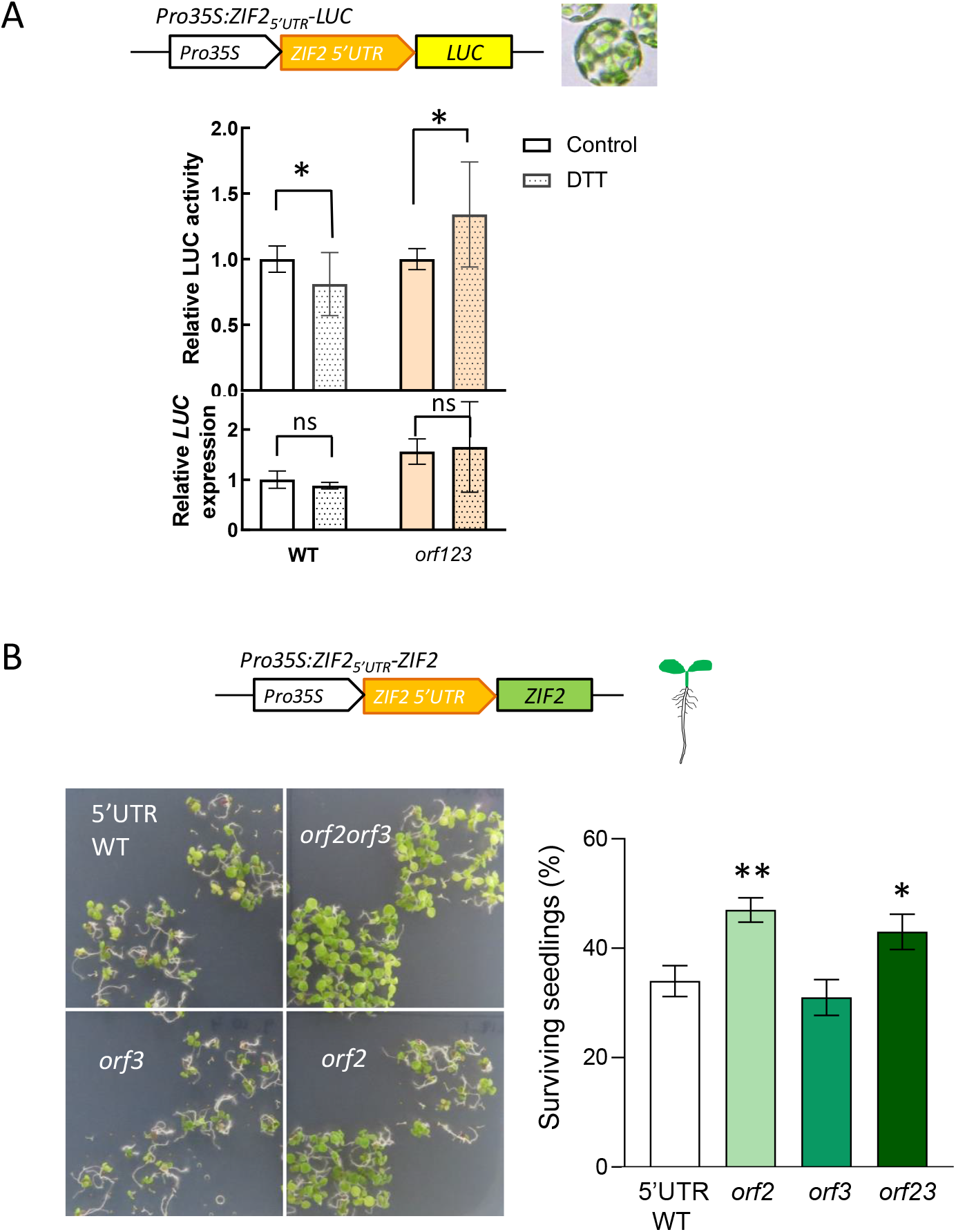
*ZIF2* uORFs role under stress conditions. A. Effect on luciferase activity of Zn and DTT treatments in mesophyll *Arabidopsis* protoplasts, comparing *ZIF2* WT 5’UTR and ZIF2 5’UTR where all the uORFs are mutated in the start codon. Bars indicate average of 5 independent experiments ± SE. B. Effect of DTT on the survival of *Arabidopsis* seedlings with *ZIF2* 5’UTR WT or mutated in the start codon of the uORFs. Bars indicate average of 5 independent experiments ± SE. Asterisks indicate significant differences with WT.

### Physiological relevance of uORFs-mediated inhibition in response to stress

uORFs are known to have a prominent role in translational inhibition under stress conditions (Son & Park 2023). The ZIF2 membrane protein has been reported to function as a zinc transporter in *A. thaliana* (Remy et al 2014). In response to a zinc challenge, total *ZIF2* expression is upregulated and the efficiency of translation increased by favoring the expression of a splice variant with an intron retained in the 5’ UTR. To gain insight into the physiological relevance of the *ZIF2* uORFs in the response to stress, their effect on translation of the encoded membrane transporter was analyzed under different stress conditions. We first tested the potential role of *ZIF2* uORF in response to Zn stress by incubating protoplasts either in control solution or with Zn, in two different constructs, WT 5’UTR or no uORFs in the 5’UTR. Zn increased translation in the WT construct, but to the same extent than in the no uORFs construct, meaning that *ZIF2* uORFs do not have a role in response to Zn stress (data not shown).

Since heavy metal stress can lead to ER stress and misfolding of proteins (De Benedictis 2023, Demircan et al 2024) and considering the nature of ZIF2 substrate, we then incubated *Arabidopsis* protoplasts with DTT, which causes ER stress. Results show that DTT decreases LUC activity in the 5’UTR WT construct, but this pattern changes when the start codon of all *ZIF2* uORFs are mutated, suggesting a role for *ZIF2* uORFs in the cellular response to ER stress. To test this hypothesis, we took advantage of the plants we generated with mutated start codons of uORF2 and uORF3 (Fig. 6). We germinated seeds of the different genotypes in DTT-containing MS plates. DTT causes an impairment in seedling development, more visible at 14 days after sowing. This effect is stronger in 5’UTR WT and *orf3* plants. However, *orf2* and *orf23* plants show a higher rate of survival after a prolonged DTT exposure, suggesting an unexplored role for ZIF2 in ER stress, mediated by translational regulation. In this way, the presence of ORF2 keeps a lower rate of ZIF2 protein, but when ORF2 is not translated (*orf2* plants), ZIF2 increases its levels conferring tolerance to DTT-triggered ER stress.

## DISCUSSION

### *ZIF2* uORF2 and uORF3 inhibit mORF translation through a fail-safe mechanism

In recent years, uORFs have arisen as key elements in the regulation of protein translation, with a promising future in engineering plants to cope with stress without fitness costs (Xu et al 2017, Zhang et al 2018) and enhancing desirable crop traits (Xing et al 2020, Yang et al 2025). However, the amount of well-characterized translated uORFs is still scarce. Here, we demonstrated that ZIF2, a zinc transporter, shows uORF-mediated translational regulation. Out of the three uORFs in *ZIF2* 5’UTR, we showed that uORF1 is not translated, thereby confirming its lack of regulatory function, while uORF2 and uORF3 act as negative regulators of the main open reading frame (mORF). This regulatory system was proved not only *in vivo* in mesophyll *Arabidopsis* protoplasts (Fig. 2C) but also *in planta* (Fig. 6).

Moreover, we report that in the absence of the constitutively active uORF2, uORF3 is translated and repress the translation of the mORF, pointing out the critical importance of this mechanism of translational regulation. To the best of our knowledge, this is the first time a fail-safe mechanism in translational inhibition has been described in plants. However, a fail-safe mechanism has been previously described in other non-plant species. The gene *GCN4* of *Saccharomyces cerevisiae* encodes 4 uORF in its mRNA, two of which cooperate in a fail-safe fashion to respond to stress (Gunisova & Valasek 2014). The mammalian orthologue of this gene, *ATF4*, encodes as well uORFs exerting a fail-safe response (Vattem & Wek 2004). This fail-safe mechanism allows cells to respond rapidly and selectively to stress by reprogramming translation, even when global protein synthesis is downregulated. Under normal conditions, ribosomes translate uORF1 and reinitiates in uORF2 (which overlaps *ATF4* mORF, thus suppressing translation of the mORF, keeping low levels of the *ATF4* protein. Under nutrient depletion, eIF2α is phosphorylated and there is less ternary complex available, translating uORF1 but bypassing uORF2, hence allowing translation of mORF, even when global translation is decreased, highlighting the importance of this protein in stress conditions. In a similar fashion, *ZIF2* uORF2 absence allows uORF3 to translate, maintaining the repression over ZIF2 translation, keeping ZIF2 at lower levels.

### uORF2 inhibits translation likely through ribosome stalling

Although uORFs are features very common in genomes (35% of genes in *A. thaliana*), the prevalence is different depending on the gene function. For instance, housekeeping genes have usually short 5’UTRs with none or few uORFs, but transcription factors and other low-expressed mRNAs usually present long 5’UTRs with abundant uORFs. Transporters have not arisen as a group enriched in uORFs, but so far, three transporters (NIP5;1, BOR1, PHO1) have already been described to have translational regulation mediated by uORFs (Tanaka et al 2016, Aibara et al 2018, Reis et al 2020). Strikingly, two of them use Boron as their substrate, but their mechanism of action differ: while in *NIP5;1* translational regulation relies on ribosome stalling *via* stabilization of eukaryotic release factor 1 in the A-site of the ribosome and impeding the reuse of 80S ribosome for downstream translation (Tanaka et al 2016, 2024), B-dependent translation of *BOR1* is mediated by translational reinitiation (Aibara et al 2018).

In the case of *ZIF2* uORFs, through a combination of targeted mutagenesis, luciferase reporter assays, and co-transfection experiments, we systematically ruled out alternative explanations (nonsense-mediated mRNA decay (NMD), *trans* effect) and demonstrated that *ZIF2* uORFs exert their inhibitory function through ribosome stalling (Figs 4,5). Ribosome stalling is not a rare translational control mechanism, but rather a recurring regulatory feature (Rhamani et al 2009, Ribone et al 2017, Sotta et al 2021). The best-characterized ribosome stalling mechanism in plants involves the nascent peptide encoded by the uORF itself. Additionally, some plant uORFs act as metabolite sensors, where ribosome stalling depends on small molecules, such as polyamines (van der Horst et al 2019).

It has been suggested that ribosome stalling can be caused by the interaction of sequence-dependent nascent peptides with some components of the exit tunnel of the ribosome (von Arnim et al 2014). Although the majority of uORFs modulate the translation of the mORF independently of their sequence, some uORFs show regulatory effects through mechanisms that rely on the peptide sequence. uORFs that exert their function in a sequence-dependent manner have been reported to belong to the so-called conserved peptide uORFs (CPuORFs). Bioinformatic analysis identified 81 highly conserved CPuORF homology groups that are present in plants, with potentially many additional groups. A considerable fraction of these may be responsive to cofactors such as sucrose or polyamines (van der Horst et al 2019). Nevertheless, although the *ZIF2* uORFs do not fall into any CPuORF category, uORF2 mediate translational control of the mORF in a sequence-dependent manner (Fig. 5C), thus providing the first evidence of a plant uORF whose regulatory function is peptide sequence–dependent without being classified as a CPuORF. It was hypothesized that uORFs with highly conserved C-terminal sequences would preferentially promote ribosome stalling *via* interactions of the nascent peptide with specific constituents of the ribosomal exit tunnel (Ebina et al 2015). When searching for critical amino-acids in *ZIF2* uORF-mediated translational regulation, an Ala scanning mutagenesis revealed that G7H8K9 residues, located in the central part of the translated peptide, have a main role on uORF2 regulatory action (Fig. 5G).

On the other hand, the functionality of uORF3 does not seem to depend on its amino acid sequence (Fig. 5C), nor on the context of the start codon (Fig. S1), which is particularly important in sequence-independent uORFs. Having discarded other mechanisms (Figs. 4, 5), uORF3 mode of action remains elusive.

### Disruption of uORF2 increases tolerance to endoplasmic reticulum stress in plants

Although numerous studies over the past two decades have described the inhibitory effects of individual uORFs, only a limited number have demonstrated their ability to repress mORF translation *in planta* (Ribone et al 2017, Aibara et al 2018, Reis et al 2020). In the case of *ZIF2* uORF2 and uORF3, we were able to demonstrate that plants carrying a mutated version of uORF2, alone or in combination with uORF3, produced more ZIF2-YFP protein than the 5’UTR WT version (Fig. 6). However, WT and mutated plants did not show differences in development, which is not surprising, given that most uORFs have shown a phenotype under stress environments.

Molecular adaptation to stress encompasses a wide range of complex responses that involve changes in gene expression, including translation. All stages of translation are tightly regulated, with initiation representing the main point of translational control. Under stress conditions, global translation is generally inhibited. However, a subset of mRNAs is efficiently translated during stress, thereby enabling plant adaptation to these conditions. Interestingly, many of these transcripts that are translated under stress harbor uORFs in their UTRs that regulate translation in response to stress conditions (Liu & Qian 2014, Young & Wek 2016, Wu et al 2024). Considering that most stress-responsive transcripts that contain uORFs are activated in response to specific environmental conditions, we analyzed whether zinc caused a translational regulation mediated by uORFs. Unfortunately, no statistical differences were found in LUC activity when incubating with ZnSO_4_ *Arabidopsis* protoplasts, transfected with a construct harboring all three uORFS in the 5’UTR *vs* all uORFs mutated (data not shown).

On the other hand, coping with severe stresses requires the synthesis and folding of a plethora of proteins in order to protect cells from the negative effects of stress. It is well stablished that in response to different stresses (such as high salinity, drought and high temperatures), the high demand of newly synthetized proteins overloads the folding capacity of the endoplasmic reticulum ER, leading to the accumulation of unfolded proteins and causing what is known as ER stress (Deng et al 2013, Liu et al 2016). Recently, it has been postulated that some heavy metals such as Cd, Zn, Cu, As can cause ER stress (De Benedictis 2023, Demircan et al 2024). In light of this, we tested the possibility of the uORF-mediated translational regulation of ZIF2, being a Zn transporter, could have a role under ER stress. For this, we triggered ER stress with DTT and our results show that plants with *ZIF2* uORFs abolished (by mutating their start codons) perform better under ER stress. The rate of survival for both *orf2* and *orf23* was higher and the plants looked healthier than their counterparts 5’UTR WT and *orf3*. We suggest that, in control conditions, uORF2 is translated by the ribosome, keeping a lower level of ZIF2 protein. However, in the absence of uORF2 (not being translated), ZIF2 protein increases favoring plant survival under ER conditions. The fine-tuning of translational regulation of the ZIF2 transporter uncovered in this work sheds new insight on the mechanisms governing plant responses to abiotic stress.

## AUTHOR CONTRIBUTIONS

EN-U and PD conceived the project and designed the research. EN-U and DS performed the experiments. EN-U and PD analyzed the data and wrote the manuscript. All authors contributed to the interpretation of results, critically reviewed the manuscript, and approved its final version.

## ACKNOWLEDGMENTS

We thank L. Romão for numerous productive discussions and V. Nunes for excellent plant care. This work was supported by the Fundação para a Ciência e a Tecnologia (FCT) through postdoctoral fellowship SFRH/BPD/112587/2015 and grant PTDC/ASP-PLA/6105/2020, as well as by the Comunidad de Madrid Atracción de Talento Investigador program (grant 2020-T1/BIO-19900). Funding from the research unit GREEN-it Bioresources for Sustainability (UIDB/04551/2020) is also acknowledged.

## SUPPLEMENTARY FIGURES

**Figure S1.** AUG context is not important for uORF2- and uORF3-mediated inhibition of mORF translation. Strong Kozak sequences of uORF2 and uORF3 were mutated to create a weaker context for translation, promoting leaking scanning of the ribosome. Effect on luciferase activity of mutating *ZIF2* uORF2 and uORF3 to weaker Kozak sequences and promote leaky scanning. Average ± SD; *n* = 4. Different letters indicate statistically significant differences (ANOVA).

**Figure S2.** Determination by quantitative RT-PCR of *LUC* transcript levels after co-transfection of *Arabidopsis* protoplasts with LUC vectors of different origins, firefly (*Photinus pyralis*, F-LUC) and sea pansy (*Renilla reniformis*, R-LUC), compared to *GUS* transcript levels. Constructs derived from each species harboring the *ZIF2* 5’UTR with only one uORF active were compared with the vector harboring all uORFs mutated. Bars indicate average of 4 independent experiments ± SE. No statistical differences were found (ANOVA).

## References

Aibara I, Hirai T, Kasai K, Takano J, Onouchi H, Naito S, Fujiwara T, Miwa K. Boron-Dependent Translational Suppression of the Borate Exporter BOR1 Contributes to the Avoidance of Boron Toxicity. Plant Physiol. 2018. 177(2):759–774. doi: 10.1104/pp.18.00119.

Baird TD, Palam LR, Fusakio ME, Willy JA, Davis CM, McClintick JN, Anthony TG, Wek RC. Selective mRNA translation during eIF2 phosphorylation induces expression of IBTKα. Mol Biol Cell. 2014 May;25(10):1686–97. doi: 10.1091/mbc.E14-02-0704

Calvo SE, Pagliarini DJ, Mootha VK. Upstream open reading frames cause widespread reduction of protein expression and are polymorphic among humans. Proc Natl Acad Sci U S A. 2009 May 5;106(18):7507–12. doi: 10.1073/pnas.0810916106.

Chang KS, Lee SH, Hwang SB, Park KY. Characterization and translational regulation of the arginine decarboxylase gene in carnation (Dianthus caryophyllus L.). Plant J. 2000 Oct;24(1):45–56. doi: 10.1046/j.0960-7412.2000.00854.x.

Clough SJ, Bent AF. Floral dip: a simplified method for Agrobacterium-mediated transformation of Arabidopsis thaliana. Plant J. 1998. 16:735–743. doi: 10.1046/j.1365-313x.1998.00343.x.

Combier JP, de Billy F, Gamas P, Niebel A, Rivas S. Trans-regulation of the expression of the transcription factor MtHAP2-1 by a uORF controls root nodule development. Genes Dev. 2008 Jun 1;22(11):1549–59. doi: 10.1101/gad.461808.

Confraria A, Baena-González E. Using Arabidopsis Protoplasts to Study Cellular Responses to Environmental Stress. Methods Mol Biol. 2016;1398:247–69. doi: 10.1007/978-1-4939-3356-3_20.

De Benedictis M, Gallo A, Migoni D, Papadia P, Roversi P, Santino A. Cadmium treatment induces endoplasmic reticulum stress and unfolded protein response in Arabidopsisthaliana. Plant Physiol Biochem. 2023 Mar;196:281–290. doi: 10.1016/j.plaphy.2023.01.056.

Demircan N, Ozgur R, Turkan I, Uzilday B. Heavy metal toxicity leads to accumulation of insoluble proteins and induces endoplasmic reticulum stress-specific unfolded protein response in Arabidopsis thaliana. Environ Sci Pollut Res Int. 2024 Aug;31(40):53206–53218. doi: 10.1007/s11356-024-34780-y.

Deng Y, Srivastava R Howell SH. Endoplasmic reticulum (ER) stress response and its physiological roles in plants. International Journal of Molecular Sciences 2013. 14(4): p. 8188–212. doi: 10.3390/ijms14048188.

Ebina I, Takemoto-Tsutsumi M, Watanabe S, Koyama H, Endo Y, Kimata K, Igarashi T, Murakami K, Kudo R, Ohsumi A, Noh AL, Takahashi H, Naito S, Onouchi H. Identification of novel Arabidopsis thaliana upstream open reading frames that control expression of the main coding sequences in a peptide sequence-dependent manner. Nucleic Acids Res. 2015 Feb 18;43(3):1562–76. doi: 10.1093/nar/gkv018.

Gunisova S, Valasek LS. Fail-safe mechanism of GCN4 translational control—uORF2 promotes reinitiation by analogous mechanism to uORF1 and thus secures its key role in GCN4 expression. Nucleic Acids Res 2014;42(9):5880–93.

Gurr SJ, Rushton PJ. Engineering plants with increased disease resistance: how are we going to express it? Trends Biotechnol. 2005 Jun;23(6):283–90. doi: 10.1016/j.tibtech.2005.04.009

Hanfrey C, Franceschetti M, Mayer MJ, Illingworth C, Michael AJ. Abrogation of upstream open reading frame-mediated translational control of a plant S-adenosylmethionine decarboxylase results in polyamine disruption and growth perturbations. J Biol Chem. 2002 Nov 15;277(46):44131–9. doi: 10.1074/jbc.M206161200

Hu Q, Merchante C, Stepanova AN, Alonso JM, Heber S. Genome-Wide Search for Translated Upstream Open Reading Frames in Arabidopsis Thaliana. IEEE Trans Nanobioscience. 2016 Mar;15(2):148–57. doi: 10.1109/TNB.2016.2516950

Kim BH, Cai X, Vaughn JN, von Arnim AG. On the functions of the h subunit of eukaryotic initiation factor 3 in late stages of translation initiation. Genome Biol. 2007;8(4):R60. doi: 10.1186/gb-2007-8-4-r60.

Kovtun Y, Chiu WL, Zeng W, Sheen J. Suppression of auxin signal transduction by a MAPK cascade in higher plants. Nature. 1998. 395:716–720. doi: 10.1038/27240

Kozak M. Effects of intercistronic length on the efficiency of reinitiation by eucaryotic ribosomes. Mol Cell Biol. 1987 Oct;7(10):3438–45. doi: 10.1128/mcb.7.10.3438-3445

Liu & Qian. WIREs RNA 2014, 5:301–315. doi: 10.1002/wrna.1212

Liu JX, Howell SH. Managing the protein folding demands in the endoplasmic reticulum of plants. New Phytol. 2016 Jul;211(2):418–28. doi: 10.1111/nph.13915

Livak KJ, Schmittgen TD. Analysis of relative gene expression data using real-time quantitative PCR and the 2(-Delta Delta C(T)) Method. Methods. 2001 Dec;25(4):402–8. doi: 10.1006/meth.2001.1262

Martinho C, Confraria A, Elias CA, Crozet P, Rubio-Somoza I, Weigel D, Baena-González E. Dissection of miRNA pathways using arabidopsis mesophyll protoplasts. Mol Plant. 2015 Feb;8(2):261–75. doi: 10.1016/j.molp.2014.10.003

Niño-González M, Novo-Uzal E, Richardson DN, Barros PM, Duque P. More Transporters, More Substrates: The Arabidopsis Major Facilitator Superfamily Revisited. Mol Plant. 2019 Sep 2;12(9):1182–1202. doi: 10.1016/j.molp.2019.07.003.

Rahmani F, Hummel M, Schuurmans J, Wiese-Klinkenberg A, Smeekens S, Hanson J. Sucrose control of translation mediated by an upstream open reading frame-encoded peptide. Plant Physiol. 2009 Jul;150(3):1356–67. doi: 10.1104/pp.109.136036.

Reis RS, Deforges J, Sokoloff T, Poirier Y. Modulation of Shoot Phosphate Level and Growth by PHOSPHATE1 Upstream Open Reading Frame. Plant Physiol. 2020 Jul;183(3):1145–1156. doi: 10.1104/pp.19.01549.

Remy E, Cabrito TR, Batista RA, Hussein MA, Teixeira MC, Athanasiadis A, Sá-Correia I, Duque P. Intron retention in the 5’UTR of the novel ZIF2 transporter enhances translation to promote zinc tolerance in arabidopsis. PLoS Genet. 2014 May 15;10(5):e1004375. doi: 10.1371/journal.pgen.1004375.

Ribone PA, Capella M, Arce AL, Chan RL. A uORF Represses the Transcription Factor AtHB1 in Aerial Tissues to Avoid a Deleterious Phenotype. Plant Physiol. 2017 Nov;175(3):1238–1253. doi: 10.1104/pp.17.01060.

Son S, Park SR. Plant translational reprogramming for stress resilience. Front Plant Sci. 2023 Feb 24;14:1151587. doi: 10.3389/fpls.2023.1151587

Sotta N, Chiba Y, Miwa K, Takamatsu S, Tanaka M, Yamashita Y, Naito S, Fujiwara T. Global analysis of boron-induced ribosome stalling reveals its effects on translation termination and unique regulation by AUG-stops in Arabidopsis shoots. Plant J. 2021 Jun;106(5):1455–1467. doi: 10.1111/tpj.15248

Tanaka M, Sotta N, Yamazumi Y, Yamashita Y, Miwa K, Murota K, Chiba Y, Hirai MY, Akiyama T, Onouchi H, Naito S, Fujiwara T. The Minimum Open Reading Frame, AUG-Stop, Induces Boron-Dependent Ribosome Stalling and mRNA Degradation. Plant Cell. 2016 Nov;28(11):2830–2849. doi: 10.1105/tpc.16.00481.

Tanaka M, Yokoyama T, Saito H, Nishimoto M, Tsuda K, Sotta N, Shigematsu H, Shirouzu M, Iwasaki S, Ito T, Fujiwara T. Boric acid intercepts 80S ribosome migration from AUG-stop by stabilizing eRF1. Nat Chem Biol. 2024 May;20(5):605–614. doi: 10.1038/s41589-023-01513-0

van der Horst S, Snel B, Hanson J, Smeekens S. Novel pipeline identifies new upstream ORFs and non-AUG initiating main ORFs with conserved amino acid sequences in the 5’ leader of mRNAs in Arabidopsis thaliana. RNA. 2019 Mar;25(3):292–304. doi: 10.1261/rna.067983.118.

Vattem KM, Wek RC. Reinitiation involving upstream ORFs regulates ATF4 mRNA translation in mammalian cells. Proc Natl Acad Sci U S A. 2004 Aug 3;101(31):11269–74. doi: 10.1073/pnas.0400541101

von Arnim AG, Jia Q, Vaughn JN. Regulation of plant translation by upstream open reading frames. Plant Sci. 2014 Jan;214:1–12. doi: 10.1016/j.plantsci.2013.09.006.

Wu HL, Jen J, Hsu PY. What, where, and how: Regulation of translation and the translational landscape in plants. Plant Cell. 2024 May 1;36(5):1540–1564. doi: 10.1093/plcell/koad197.

Xing S, Chen K, Zhu H, Zhang R, Zhang H, Li B, Gao C. Fine-tuning sugar content in strawberry. Genome Biol. 2020 Sep 3;21(1):230. doi: 10.1186/s13059-020-02146-5.

Xu G, Yuan M, Ai C, Liu L, Zhuang E, Karapetyan S, Wang S, Dong X. uORF-mediated translation allows engineered plant disease resistance without fitness costs. Nature. 2017 May 25;545(7655):491–494. doi: 10.1038/nature22372

Yang Q, Zhu W, Tang X, Wu Y, Liu G, Zhao D, Liu Q, Zhang Y, Zhang T. Improving rice grain shape through upstream ORF editing-mediated translation regulation. Plant Physiol. 2024 Dec 23;197(1):kiae557. doi: 10.1093/plphys/kiae557

Yoo SD, Cho YH, Sheen J. Arabidopsis mesophyll protoplasts: a versatile cell system for transient gene expression analysis. Nat Protoc. 2007;2(7):1565–72. doi: 10.1038/nprot.2007.199.

Young SK, Wek RC. Upstream Open Reading Frames Differentially Regulate Gene-specific Translation in the Integrated Stress Response. J Biol Chem. 2016 Aug 12;291(33):16927–35. doi: 10.1074/jbc.R116.733899

Zhang H, Si X, Ji X, Fan R, Liu J, Chen K, Wang D, Gao C. Genome editing of upstream open reading frames enables translational control in plants. Nat Biotechnol. 2018 Oct;36(9):894–898. doi: 10.1038/nbt.4202.

Zhang T, Wu A, Yue Y, Zhao Y. uORFs: Important Cis-Regulatory Elements in Plants. Int J Mol Sci. 2020 Aug 28;21(17):6238. doi: 10.3390/ijms21176238

