## Supplementary figures for "Upstream ORFs regulate translation of the Arabidopsis ZIF2 transporter and affect ER stress tolerance"

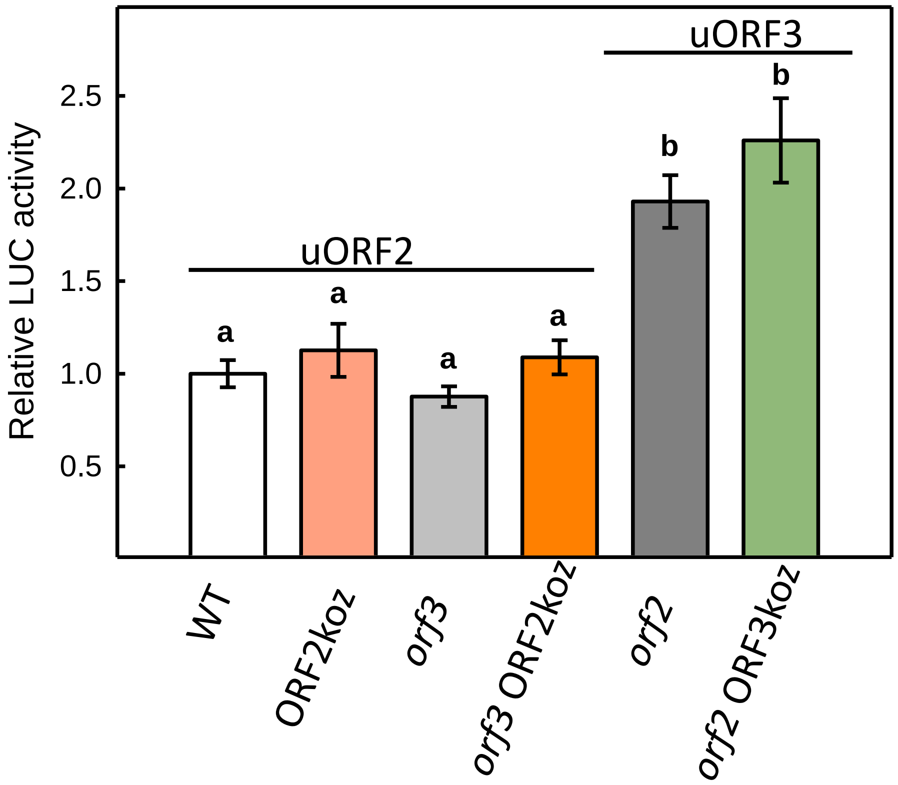


**Figure S1.** AUG context is not important for uORF2- and uORF3- mediated inhibition of mORF translation. Strong Kozak sequences of uORF2 and uORF3 were mutated to create a weaker context for translation, promoting leaking scanning of the ribosome. Effect on luciferase activity of mutating *ZIF2* uORF2 and uORF3 to weaker Kozak sequences and promote leaky scanning. Average ± SD; *n* = 4. Different letters indicate statistically significant differences (ANOVA).

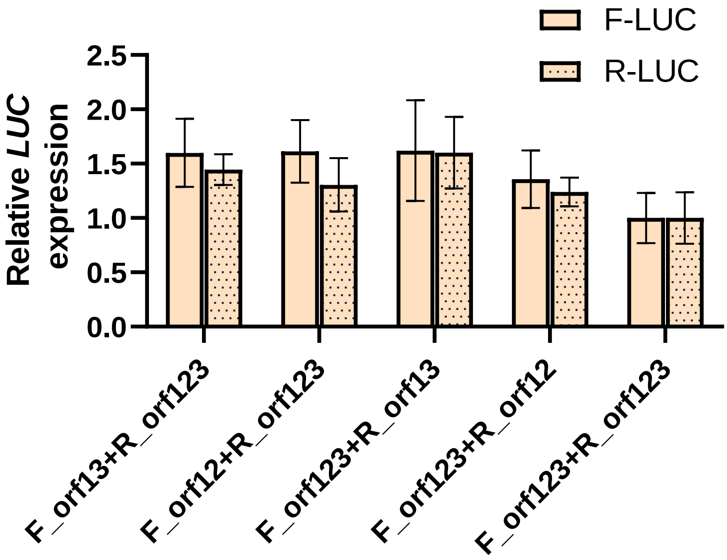


**Figure S2.** Determination by quantitative RT-PCR of *LUC* transcript levels after co-transfection of arabidopsis protoplasts with LUC vectors of different origins, firefly (*Photinus pyralis,* F-LUC) and sea pansy (*Renilla reniformis,* R-LUC), compared to *GUS* transcript levels. Constructs derived from each species harboring the *ZIF2* 5'UTR with only one uORF active were compared with the vector harboring all uORFs mutated. Bars indicate average of 4 independent experiments ± SE. No statistical differences were found (ANOVA).
